# CCK* (Convex Closure K*): A Suite of Algorithms for De Novo L- and D-peptide Design

**DOI:** 10.1101/2025.11.21.689740

**Authors:** Henry Childs, Allen C. McBride, Bruce R. Donald

**Affiliations:** Department of Chemistry, Duke University, 124 Science Drive, 27708, North Carolina, USA; Department of Computer Science, Duke University, 308 Research Drive, 27708, North Carolina, USA; Department of Biochemistry, Duke University School of Medicine, 307 Research Drive, 27710, North Carolina, USA; Department of Mathematics, Duke University, 120 Science Drive, 27708, North Carolina, USA

**Keywords:** algorithms, protein structure, OSPREY, D-peptides, structure-based drug design, computational biology

## Abstract

The computational design of L-peptides and their mirror-image counterparts, D-peptides, is an active area in drug design. Peptide therapeutics offer exceptional structural diversity and high binding specificity, while D-peptides additionally confer critical advantages such as proteolytic resistance. Progress in de novo D-peptide design has been hindered by the absence of evolutionary context and limited structural data, both of which underpin the deep learning methods widely used in L-peptide design. Consequently, a robust framework capable of designing both L- and D-peptides should integrate data-driven inference with first-principles, physics-based modeling. Here, we introduce a unified computational framework that supports de novo design of both L- and D-peptides, thereby expanding the accessible design space across both chiral spaces. Convex Closure K* (CCK*) is a suite of chirality-agnostic algorithms: SCOPE, MONTAGE, and ARISE. SCOPE uses geometry as a proxy for chemical energetics, computing convex hull representations of rotameric states to rapidly generate multi-sequence protein contact maps. MONTAGE employs geometric hashing in conjunction with the K* algorithm to generate and rank backbone scaffolds according to their suitability for sequence design. ARISE is a K*-based sequence design algorithm that performs iterative residue assignment in an undirected graph to design high-affinity peptide sequences. We apply the full CCK* suite to six de novo design tasks, benchmarking chirality-preserving and chirality-inverting designs in both homochiral and heterochiral complexes.

## Introduction

Relative to small molecules and biologics, peptide therapeutics demonstrate advantages including high specificity, enhanced membrane permeability, and improved bioavailability (1). D-peptides, the mirror images of proteinogenic L-peptides, resist proteolytic degradation and thus exhibit improved stability (2), among other benefits. Due to these favorable properties, L- and D-peptides are promising modalities for drug design.

Peptides are difficult to design *in silico*. Discrete rotamer optimization for identifying the global minimum energy conformation (GMEC) is NP-hard (3). Deep learning methods show promise for L-peptide binder design (4, 5) and have been extended to systems containing noncanonical amino acid substitutions (6, 7). However, likely due to the scarcity of experimental structures and absence of evolutionary data, these methods can fail to predict D-peptide binders (8). Thus, a generalizable framework for designing L- and D-peptides must remain effective in a data-sparse regime. An algorithmic framework capable of *de novo* L- and D-peptide design must address three fundamental challenges: (1) generating and ranking realistic candidate binding poses, (2) tractably exploring conformation and sequence space, and (3) accurately predicting binding affinity. Herein, we review prior work addressing these challenges.

Several previous strategies for D-peptide design have focused on sampling from experimentally determined structures to generate a scaffold library. Garton et al. (9) searched a mirror image of the PDB and identified hotspot residues to design D-peptide GLP1R and PTH1R agonists. We previously reported on DexDesign (10, 11), an algorithm for *de novo* D-peptide design that uses reflections and geometric hashing to generate a scaffold library. D-peptides produced by DexDesign were predicted to bind disease-associated PDZ domains. Experimental techniques for identifying D-peptide binders often rely on screening a large phage display library (12, 13).

Physics-based methods have a strong track record in sequence and conformation optimization. Methods implementing deterministic search algorithms, such as dead-end elimination (14, 15), generate predictions with provable guarantees on accuracy relative to the energy model. These methods have been applied to diverse chemical systems such as HIV-1 V2-apex antibody redesigns (16) and KRas binder optimization (17). Similar approaches have been applied to non-proteinogenic systems (10, 18). Rosetta (19, 20) incorporates D-amino acid templates for noncanonical design; these methods rely on stochastic sampling rather than deterministic optimization with formal guarantees. Cost function networks have also been applied for deterministic discrete optimization in protein design (21), but, to our knowledge, have not been applied to D-peptide design.

The thermodynamic calculation of the association constant (*K*_*a*_) employs a partition function (*Z*) for the apo protein (*P*), apo ligand (*L*), and protein:ligand complex (*PL*) (22):

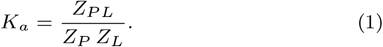

In this formulation, calculation of binding affinity relies on accurate partition functions. Accurate partition functions, in turn, depend on conformational ensembles that adequately sample the relevant regions of the energy landscape to capture the entropic contributions governing protein interactions. Physics-based docking methods (23) and deep learning models (24) can generate ensembles. Although these approaches are enticing for enabling broad searches over large-scale conformational changes, they generally do not offer formal guarantees that low-energy states are represented. Because Boltzmann weighting exponentially amplifies contributions from low-energy states, failure to include relevant energy minima when calculating the partition function compromises the calculation of binding affinity.

The *K*\* algorithm and its extensions (25–30) bound the required partition functions with provable guarantees of free energy approximation. For each species *i* ∈ {*P, L, PL*}, an approximation to the partition function (*Q*_*i*_ ≈ *Z*_*i*_) is computed by summing over each state (*s*):

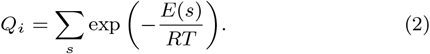

*R* and *T* are the gas constant and temperature, respectively. The minimized energy of a given state, *E*(*s*), is computed over continuous rotamers (26) for a given ensemble. *K*\* uses an AMBER-based, pairwise-decomposable molecular mechanics energy model (27, 31) with EEF1 for implicit solvation modeling (32). This energy model enables reflection operations in design by respecting energy equivariance precisely. Ensemble conformations are searched using the A* algorithm (33) and enumerated in a gap-free list ordered by lower bound on energy. By maintaining an upper bound on the partition function, unenumerated conformations are guaranteed to have higher energy (see (25) for more information). By computing an *ε*-accurate partition function for each species, *K*\* approximates the true *K*_*a*_:

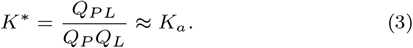

In short, *K*\* estimates binding affinity by approximating the partition functions of the unbound protein, unbound ligand, and bound complex. This yields a *K*\* score that approximates the association constant *K*_*a*_ with provable guarantees on partition function accuracy. *K*\* predictions have been shown to agree with experimental measurements, accurately ranking sequence variants by binding affinity (Spearman’s *ρ* = 0.76) (27) and quantitatively estimating ΔΔ*G*_*fold*_ (34).

Pruning high-energy conformations from consideration reduces the number of calls to the energy function, thereby accelerating *Q*_*i*_ calculation (Eq. 2) without sacrificing accuracy. Pruning algorithms offer combinatorial speedups compared to exhaustive enumeration (see multi-sequence *Q*_*i*_ bounds in *BBK*\* (28) and intra-chain pruning using Inverse Alanine Scanning (IAS) (10)); however, these algorithms are not sufficiently efficient to explore vast sequence spaces. More effective pruning is required for *de novo* design because the number of possible sequences is exponential in the number of residues. For example, IAS calculates *K*\* binding predictions for optimal point mutations on a polyalanine peptide, which is then used in a combinatorial search to construct full sequences that are predicted to bind a protein target (10). In the ideal case, IAS returns ≤ 5 non-clashing residue identities at each backbone position. A 6-residue peptide combinatorial search will need to enumerate at most 5^6^ = 15, 625 sequences of the 20^6^ possible sequences. However, in the analogous case on a 12-residue peptide, a combinatorial search must enumerate 5^12^ ≈ 244, 000, 000 sequences after IAS pruning. The pruned search space for the 12-residue peptide (5^12^ sequences) is greater than the full search space of the 6-residue peptide (20^6^ sequences). These scaling limitations motivate algorithms that can more effectively prune sequence and conformation space before exhaustive enumeration.

Here we introduce Convex Closure *K*\* (*CCK* *), a suite of generalizable algorithms (scope, montage, and arise) for the *de novo* design of L- and D-peptide binders. These algorithms extend *K*\* to yield broader applicability. We apply these algorithms in a reproducible workflow to six biological systems and validate predictions using *in silico* benchmarks. *CCK* * predicts high-affinity *de novo* L- and D-peptides spanning antitumor, antimicrobial, and CFTR-modulating drug classes. By presenting *CCK* *, this article makes the following contributions:

1. A free and open-source implementation of Convex Closure *K*\* (*CCK* *) in the OSPREY software package (27).
2. scope (Side Chain Orientation and Position Evaluation): a geometric algorithm for generating multi-sequence protein contact maps, enabling *a priori* ranking and pruning of conformation and sequence space.
3. montage (Motif-Oriented Noncanonical Template Assembly and Generation Engine): a docking and scaffold generation algorithm that uses geometric hashing and the *K*\* algorithm to return polyalanine scaffolds ranked by viability for sequence design.
4. arise (Affinity-driven Rational Iteration for Sequence Engineering): a sequence design algorithm that leverages greedy sequence assignment in an undirected graph to generate high-affinity peptide sequences.
5. Bounds on time complexity for each algorithm. Under modest assumptions, the algorithms are polynomial time.
6. Benchmarks of the *CCK* * software suite on six *de novo* design tasks spanning chirality-preserving and chirality-inverting designs in homochiral and heterochiral complexes.

## Methods

### SCOPE

In computational protein design, defining the search space can be just as important as the search itself. Calculating an accurate binding affinity (Eq. 1) relies on consideration of all relevant low-energy conformation states. Boltzmann weighting (Eq. 2) means low-energy states contribute exponentially more to affinity values; individual high-energy states are negligible. We therefore aim to identify the *productive search space* for protein design, defined as the subset of states with appreciable Boltzmann weight (i.e., states that make non-negligible contributions to the partition function). As reviewed below, models commonly implement simplifications and guiding heuristics to define the productive search space *a priori*; this strategy concentrates the search within conformational regions of maximal likely relevance.

Rotamer libraries are a common simplifying assumption in computational protein design. By extracting the modal dihedral angles from empirical rotamer populations (35), a discrete set of rotameric states can approximate the continuous search space for each proteinogenic amino acid. Reflection is an energy-equivariant geometric transformation (36), so a rotamer library for D-amino acids can be obtained via a reflection operation on L-rotamers (10). Molecular voxel theory (26) applies bounds on high-dimensional regions of configuration space (C-space) to model continuous protein flexibility for these rotamers. These representations are tractable to search using optimization algorithms (15, 33).

Protein contact maps (37, 38) are valuable for describing the spatial structure of proteins and complexes. These methods commonly use evolutionary signals from sequence homologs and probabilistic contact predictions to infer pairwise residue contacts, protein fold, and function. Deep learning methods, such as AlphaFold, use analogous pair representations in the Evoformer module (39). Other work has used linear programming over local constraints on interatomic distances to quickly prune rotamers for biophysical modeling (40).

A protein contact map for L- and D-peptide design must operate with no evolutionary priors (homochiral D-peptides are synthetic) and use novel computational strategies to accommodate the large *de novo* search space, extending contact maps beyond the traditional use in single-sequence structure inference. Here, we introduce scope (Side Chain Orientation and Position Evaluation), which extends the methodology of rotamers and protein contact maps to generate *multi-sequence protein contact maps* of protein complexes for design (see Fig. 1). By using geometry as a proxy for chemical energetics, scope rapidly defines the productive search space of all possible sequences while capturing interaction effects of side chain rotamers. scope does not require any calls to an energy function or evolutionary priors. This positions scope as a tool for rapid, large-scale exploration of design spaces in data-sparse domains: generated contact maps account for both sequence variants (mutants) and multiple side-chain rotameric states (modal images of continuous rotamers) across protein chains. scope calculates the productive search space relative to a fixed backbone and models rotamer states using convex hulls. For a given peptide:target complex, the algorithm constructs mutant hulls for peptide design positions and wild-type hulls for target residues. A mutant hull is defined as the convex polyhedron that encloses all rotamer atoms (including hydrogens) for all amino acid identities (see Fig. 1 (A)). A wildtype hull is defined as the convex polyhedron that encloses all rotamer atoms (including hydrogens) for a target residue’s wildtype identity (see Fig. 1 (B)). By representing the chemical interaction space in this manner, scope rapidly determines potential pairwise residue interactions by polyhedral C-space intersections. The productive search space is defined from the C-space as all pairs of hulls that have a volumetric overlap. The volume of polyhedral overlap (Å^3^) provides a geometric heuristic for the significance of a chemical interaction (similar to a cutoff distance (41)). Non-overlapping hulls can be inferred to have no meaningful chemical contact; this enables combinatorial pruning of sequences and conformations for design.

**Figure 1.**
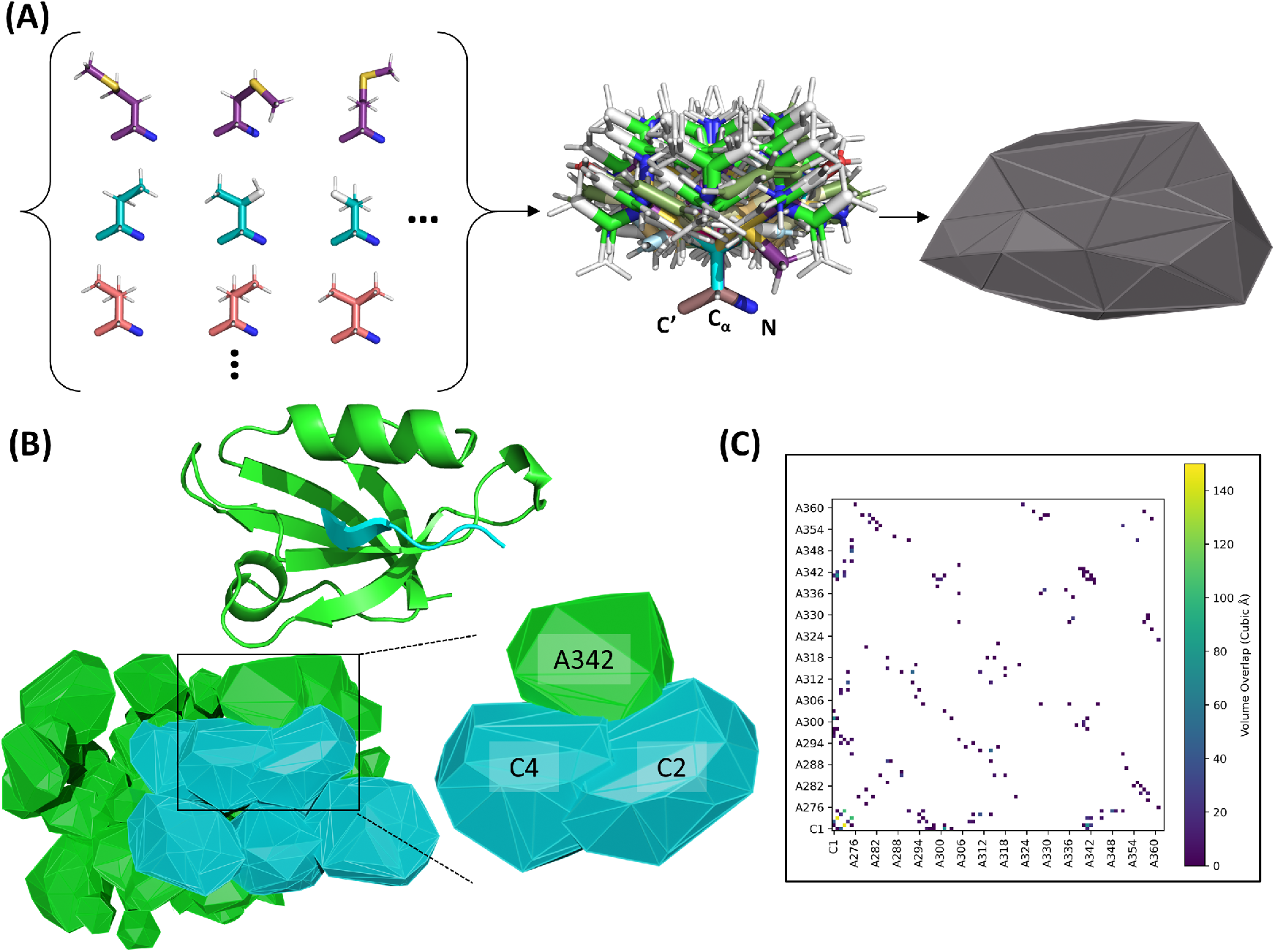
The SCOPE algorithm applied to kCAL01:CALP. **(A)** Generation of a mutant convex hull. All rotamers for the 20 L (or D) amino acids are aligned and the convex hull is computed. A wildtype hull is similarly computed using only the rotamers corresponding to the wildtype identity. These convex polyhedra represent the configuration space of amino acid side chains on a fixed backbone. **(B)** Top: Visualization of the L-peptide kCAL01 bound to the biological target CALP (PDB ID 6ov7), a validated cystic fibrosis drug target. CALP (chain A) is displayed in green; kCAL01 (chain C) in cyan. Left: the kCAL01:CALP complex represented as mutant and wildtype hulls. Right: The mutant hulls of residues C4 (wildtype Thr) and C2 (wildtype Gln) intersect with an overlap volume of 145.2 Å^3^. C4 has an overlap volume with the wildtype hull of A342 (wildtype Lys) of 13.4 Å^3^; C2 has an overlap volume of 26.3 Å^3^. **(C)** A multi-sequence protein contact map generated using scope for kCAL01:CALP.

Mutant and wildtype hulls are computed using the penultimate rotamer library (35) and an implementation of the Quickhull algorithm (42). scope is compatible with any rotamer library, but we selected the penultimate rotamer library because it has supported accurate designs in prior *K*\* work (27). The time complexity of computing a convex hull in ℝ^3^ of *p* points is 𝒪(*p* log *p*). In this work, the points are atomic coordinates. This is the most intensive computation; volume overlap operations (43) are linear in the number of points. Thus, scope has a computational complexity of 𝒪(*p* log *p*). In the systems tested here, this algorithm runs in seconds per structure.

### MONTAGE

A crucial task in protein design is generating backbone structures that are viable candidates for subsequent *sequence design*, a process in which amino acid sequences are selected to optimize desired properties such as binding affinity. Previously, our lab used the DexDesign algorithm (10) to generate a library of D-peptide candidate binders. To accomplish this, DexDesign calls MASTER (44), a geometric hashing algorithm. Given a protein structure, MASTER searches a PDB database and returns matching structures by increasing *C*_*α*_ alignment RMSD. A low RMSD value indicates close backbone agreement with the input, making this metric useful for prioritizing matches that preserve desired structural geometry. The *K*\* algorithm provides a thermodynamic measure of match quality, with higher *K*\* scores indicating more favorable starting points for sequence design. DexDesign therefore ranked design candidates by *designability*: the predicted suitability for sequence design according to *C*_*α*_ RMSD and *K*\* score. DexDesign combined geometric priors (experimentally determined structures located with MASTER) with physics-based calculations (the *K*\* algorithm (30)) to generate peptides that were predicted to bind L-targets (10). However, this algorithm relied on the wildtype sequence of the sampled structure to predict designability, which biases selection toward structures with large side chains that improve binding site packing (45). Additionally, docking was determined by geometric alignment with limited conformational sampling. This reduced the number of ensemble conformations used to calculate *K*\* scores, hindering accurate designability rankings.

MONTAGE (Motif-Oriented Noncanonical Template Assembly and Generation Engine) is an algorithm for generating L- and D-peptide scaffold libraries for protein targets. This algorithm calls scope (see Methods: SCOPE) as a subroutine to determine conformational sampling in *K*\* structural ensembles. montage mutates all ligand residues to alanine (excluding glycine and proline residues, whose unique conformational properties may be incompatible with substitution under a fixed-backbone model). This biases the *K*\* algorithm to rank scaffolds based on backbone compatibility as opposed to binding interface packing. Further, because alanine possesses only a single rotameric state, this restricts the ligand side-chain search space, enabling more sampling of target protein conformation space. Each scaffold is translated (±1.2Å) and rotated (±5^°^) during the *K*\* search to find docking configurations that better satisfy the geometric constraints of the interface. The combination of polyalanine scaffolds and improved flexibility modeling enables montage to produce more discriminative designability rankings.

Given an L-peptide:L-protein complex, montage mirrors the L-peptide to D-space and uses this structure as a query to MASTER (44). MASTER, using the L-database, returns L-substructures that are ordered by backbone similarity to the D-peptide query. Performing one final reflection operation yields D-peptide scaffolds that are geometrically similar to the endogenous L-peptide. The *K*\* algorithm is used to mutate all peptide residues (except glycine and proline) to alanine. Conformational modeling for the *K*\* calculation is defined by scope. Because MASTER returns scaffolds ordered by increasing *C*_*α*_ RMSD, subsequent ordering by *K*\* score ranks scaffolds by predicted designability. See Fig. 2 for a visualization of this methodology and SI Fig. S2 for pseudocode. L-peptide scaffolds are similarly generated by omitting reflection operations. MASTER was chosen because of prior effectiveness in D-peptide design (10); however, montage is agnostic to structure search subroutine and can use alternative tools such as Foldseek (47).

**Figure 2.**
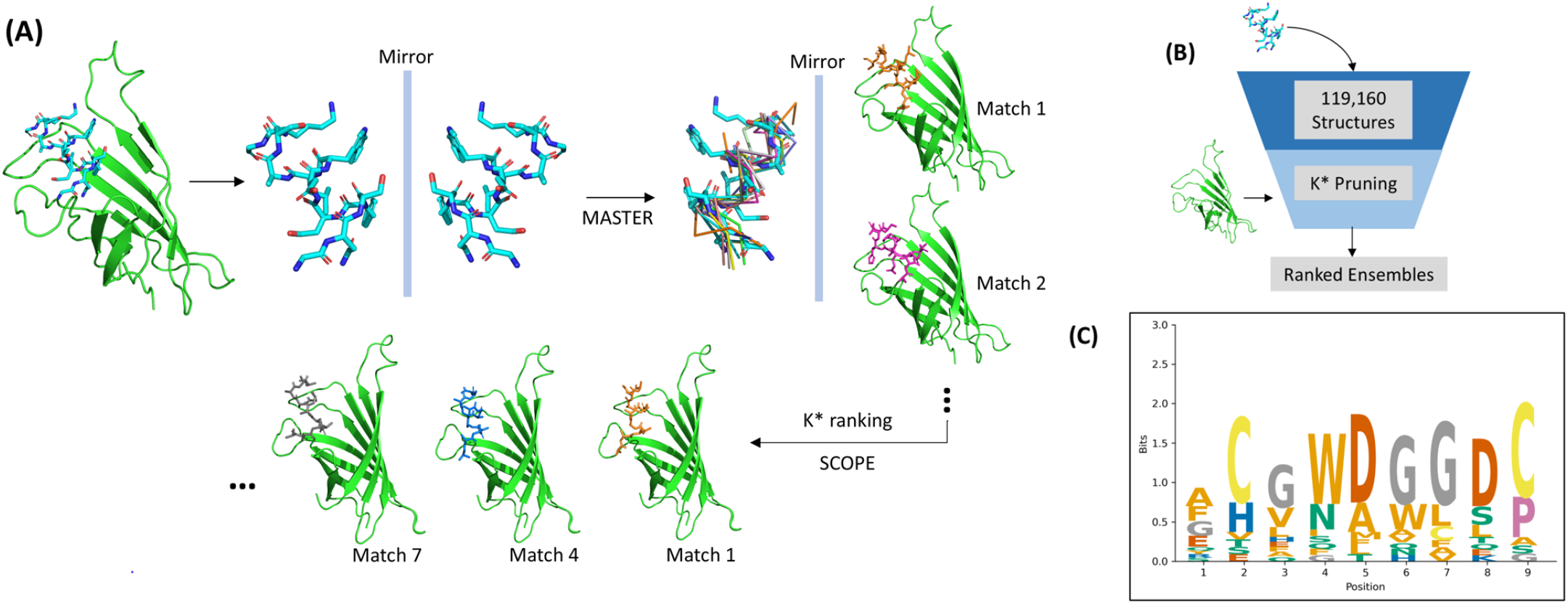
MONTAGE generates and ranks scaffolds by designability for a D-peptide:L-target system. This figure displays results for D-peptide GNSFDDWLASKG (herein referred to as GNS) bound to Streptavidin (PDB ID 5n89). The suitability of a scaffold for sequence design is referred to as designability. **(A)** The montage algorithm. Using the structure of GNS (cyan) bound to Streptavidin (green), GNS is isolated and reflected to L-space. The MASTER (44) subroutine uses geometric hashing to sample structurally similar L-backbones (multicolor sticks) from a PDB database. Matches are returned by increasing *C*_*α*_ alignment RMSD. These L-backbones are reflected back to D-space and aligned into the binding site of the target (Streptavidin), resulting in D-peptides that mimic the GNS fold. The *K*\* algorithm (25) mutates the D-scaffolds to polyalanine and ranks these complexes by computing the binding affinity over conformational ensembles. *K*\* translates and rotates the ligand in the binding site during this search. scope (see Methods: SCOPE) determines residue conformation sampling for the *K*\* search. **(B)** Overview of sequence and conformation pruning using montage. Using the structure of GNS:Streptavidin, MASTER, and the *K*\* algorithm, montage returns scaffolds ranked by designability. Our MASTER database contains 119,160 high-resolution (*≤* 2.5 Å) structures from the PDB (46). **(C)** Sequence logo for the top 20 MASTER matches of (reflected) GNS. See SI Fig. S2 for montage pseudocode.

For a non-disjoint segment with *s* residues, the time complexity for MASTER is 𝒪 (*s*^2^). MASTER computes over a million superpositions per second (44), so the empirical runtime is minutes. Conformational modeling for *K*\* calculations is determined using scope, which has a complexity of 𝒪 (*p* log *p*) (see Methods: SCOPE). The *K*\* algorithm computes binding affinity for the polyalanine scaffolds. Although *K*\* ensemble enumeration is worst-case exponential (25), branch-decomposition-based methods (41) can compute the *K*\* score (Eq. 3) in time 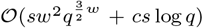. Here, *w* is branch width, *q* is the number of rotamers per residue, and *c* is the number of conformations in a partition function. For many problems, the branch width *w* is constant, reducing the complexity to polynomial time. Thus, montage has a computational complexity of 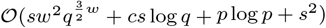.

### ARISE

In *de novo* design, the number of candidate sequences is exponential in the number of residues. This is intractable even for accelerated *K*\* algorithms such as *BBK*\* (28), which has a runtime that can be sublinear in the number of sequences. Because the sequence search space is exponentially large, the true global optimum cannot, in practice, be calculated by exhaustive enumeration using intensive physics-based methods. Nevertheless, it is possible to systematically obtain sequences that likely lie within a neighborhood of the optimum, thereby producing practical, high-scoring designs. Here, we present arise (Affinity-driven Rational Iteration for Sequence Engineering), an algorithm for *de novo* sequence selection for affinity optimization.

Given an input structure, such as a peptide:protein complex, arise constructs the undirected graph *G* = (*V, E*). In this graph, vertices (*V*) encode residues. There are two types of residues: those corresponding to design chain residues *D*, and those corresponding to target chain residues *T* (*V* = *V*_*D*_ ∪ *V*_*T*_, *V*_*D*_ ∩ *V*_*T*_ = ∅). Edges (*E*(*τ*)) represent a potential chemical interaction between vertices in *V* based on geometry; using scope (see Methods: SCOPE), an edge is drawn between two vertices if the convex hulls of those residues have an overlap greater than the volume threshold *τ*. We assign to each vertex a set of amino acid identities. Let *g* be the mapping from a vertex to a set of amino acid identities *g*: *V* → 𝒫 (*A*), where 𝒫 is the nonempty powerset and *A* is the amino acid alphabet. In effect, arise changes *g* to restrict the set of amino acids feasible at each design residue in an iterative fashion (described below). Let *c* be the mapping from a set of amino acid identities to a convex hull *c*: 𝒫 (*A*) → *S*, where *S* is the set of all convex hulls. Thus, the scope subroutine defines *E*(*τ*) using its hull volume overlap function *h*:

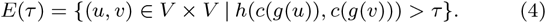

All design chain residues *v* ∈ *V*_*D*_ are initialized to *g*(*v*) = *A* and all target chain residues *v* ∈ *V*_*T*_ are initialized to their wildtype identity. Neighboring residues of a pair *v*_*i*_, *v*_*j*_ ∈ *V*_*D*_ (where (*v*_*i*_, *v*_*j*_) ∈ *E*(*τ*)) are defined:

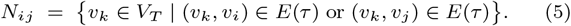

*K*^∗^((*a*_*p*_, *b*_*q*_), *M*) is defined as the binding affinity estimate obtained by running the *K*\* algorithm (Eq. 3) with modified subsequence (*a*_*p*_, *b*_*q*_) at design chain positions *p* and *q* (*p*≠ *q, a*_*p*_, *b*_*q*_ ∈ *A*). Residues in set *M* are modeled with continuous flexibility (26) during *K*\* searches. Therefore, arise computes the highest-affinity sequence pair by:

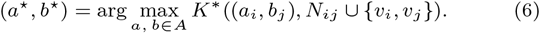

Based on Eq. 6, arise updates *g* such that *g*(*v*_*i*_) = {*a*^⋆^} and *g*(*v*_*j*_) = {*b*^⋆^}. The GMEC conformation from this *K*\* calculation is propagated as the structure for the subsequent iteration. arise calls scope to recompute edges *E*(*τ*) (Eq. 4) given these new sequence assignments. This *K*\*-based search (Eq. 6) iteratively updates *g* until all design residues have been assigned singleton amino acid identities. The *K*\* algorithm has been shown to accurately design side chain interactions in diverse chemical systems (16, 17, 45).

By modeling design chain residues as convex hulls, arise obtains an upper bound on the feasible C-space for unassigned residue identities. Each residue assignment reduces the sequence and conformation space by constricting the corresponding hull volumes, allowing re-evaluation of scope-defined interactions (Eq. 4) and edge-pruning *G*. While exhaustive exploration of the *de novo* sequence space remains intractable, this framework implements a greedy search algorithm that identifies locally optimal pairwise assignments within progressively restricted regions of sequence and conformational space.

We do not claim global optimality. This method exploits monotone pruning of *G* to remove rotamer selections that are locally suboptimal under *K*\* scoring, thereby shrinking the productive search space before subsequent sequence design. Empirically, arise does not appear susceptible to becoming trapped in local energy minima and finds similarly promising solutions across all initial conditions tested (see SI Section 1). See Fig. 3 for a visualization of this algorithm. Conceptually, this methodology is reminiscent of region-based belief propagation in factor graphs (48), but uses scope multi-sequence contact maps in place of region cutoffs.

**Figure 3.**
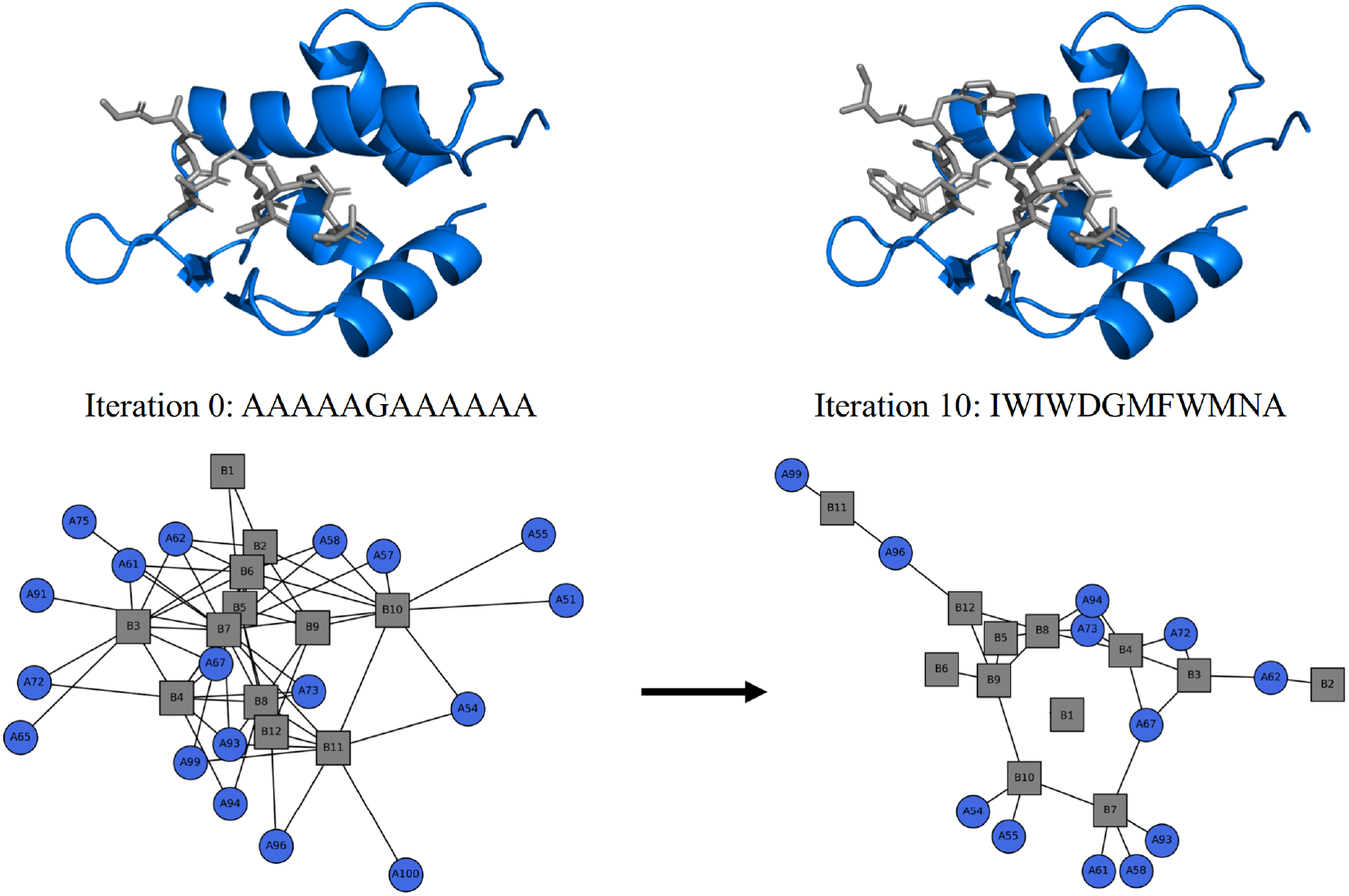
The ARISE algorithm designs a high-affinity D-peptide. This figure visualizes a montage scaffold (see Methods: MONTAGE) generated from the D-peptide:MDM2 complex (PDB ID 3iwy). **Left:** the 12-residue D-peptide scaffold (chain B) is shown in gray sticks, while L-MDM2 (chain A) is shown in blue cartoon representation. For the undirected graph generated by arise, the D-peptide residues are shown as gray boxes and MDM2 residues are shown as blue circles. Edges indicate a pairwise hull overlap exceeding the volume threshold *τ* (Eq. 4). The identity of each D-peptide scaffold residue is updated during successive *K*\* searches (Eq. 6). **Right:** the graph and designed peptide structure after ten iterations (in this case, full chain assignment). Amino acid assignment constricts design chain hull volumes, pruning graph edges and shrinking the search space for subsequent *K*\* calculations.

The arise algorithm constructs the undirected graph using scope, which has a computational complexity of 𝒪 (*p* log *p*) (see Methods: SCOPE). The iterative search is linear in the number of design residues, using the *K*\* algorithm for sequence selection. As described in Methods: MONTAGE, the computational complexity of this algorithm is 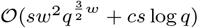.

## Results

We applied the *CCK* * software suite in a reproducible workflow (see Fig. 4(A)) to six *de novo* design tasks. We report benchmarks for homochiral (L-peptide:L-protein) and heterochiral (D-peptide:L-protein) binding. Given a structural template, we evaluated *CCK* * on both chirality-preserving designs (e.g., L-peptide template to L-peptide prediction) and chirality-inverting designs (e.g., L-peptide template to D-peptide prediction). To validate designed sequences, we investigated predicted binding affinity (*K*\* score, where a higher score indicates better binding), hydrogen bonds, and sequence recovery. All *K*\* scores are reported on a log_10_ scale. All designs were performed across five polyalanine scaffolds returned by montage. For tractability, arise *K*\* calculations were capped at 24 hours with at most two flexible target residues, selected by largest hull volume overlap with the design chain. Based on scaffold geometry, some *K*\* runs (Eq. 6) assigned only one residue, such as in cases involving an isolated vertex. Hydrogen bonds were computed from the GMEC structure using PyMOL (52). Sequence similarity was computed using the VectorBuilder sequence alignment tool (53). Each design was run on a 24-core, 48-thread Intel Xeon processor.

**Figure 4.**
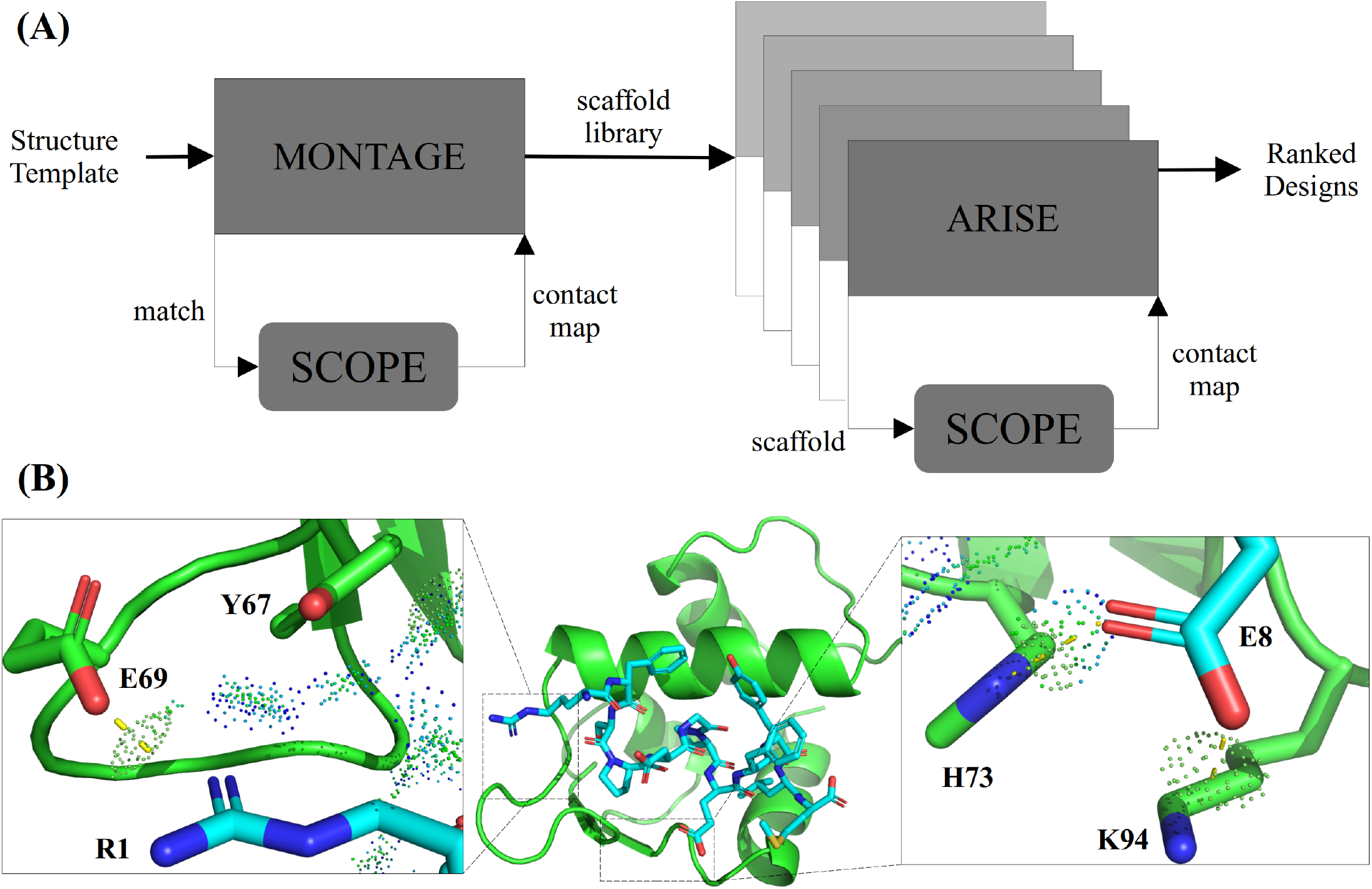
Application of the CCK* suite to a cancer-relevant system. **(A)** The *CCK* * workflow. A structural template (PDB file) of a peptide:target complex is fed into the montage algorithm (see Methods: MONTAGE). Using the template peptide, montage invokes MASTER (44) to return matching experimental structures by increasing *C*_α_ alignment RMSD. MASTER matches are aligned into the target binding interface, replacing the template peptide. The scope algorithm is invoked on each match:target complex to infer pairwise residue interactions, which determine conformational sampling in subsequent *K*\* searches. montage calls the *K*\* algorithm to mutate the MASTER matches to polyalanine, generating a ranked polyalanine scaffold library. This scaffold library is passed to arise (see Methods: ARISE). For each scaffold:target (here, shown as five complexes) arise invokes scope to construct an undirected graph representing the sequence and conformation search space. arise iteratively discovers high-affinity residues for the peptide scaffold, generating full sequences ranked by affinity. **(B)** Structural analysis of a *CCK* *-generated D-peptide inhibitor of MDM2, a p53-binding domain (49). The D-peptide is shown in cyan sticks; MDM2 is shown in green cartoon. Yellow dashes indicate inferred hydrogen bonds. Favorable van der Waals contacts are indicated by blue and green MolProbity dots (50, 51). Left: designed D-peptide residue R1 establishes an H-bond with target residue E69 and favorable vdW contacts with target residue Y67. Right: designed D-peptide residue E8 establishes H-bonds with target residues H73 and K94. Compared to known binder DP12, the designed D-peptide has a better predicted binding affinity (*K*\* score = +6.6, ΔΔ*G ≈ −*9 kcal/mol) and establishes one more hydrogen bond with MDM2.

**Figure 5.**
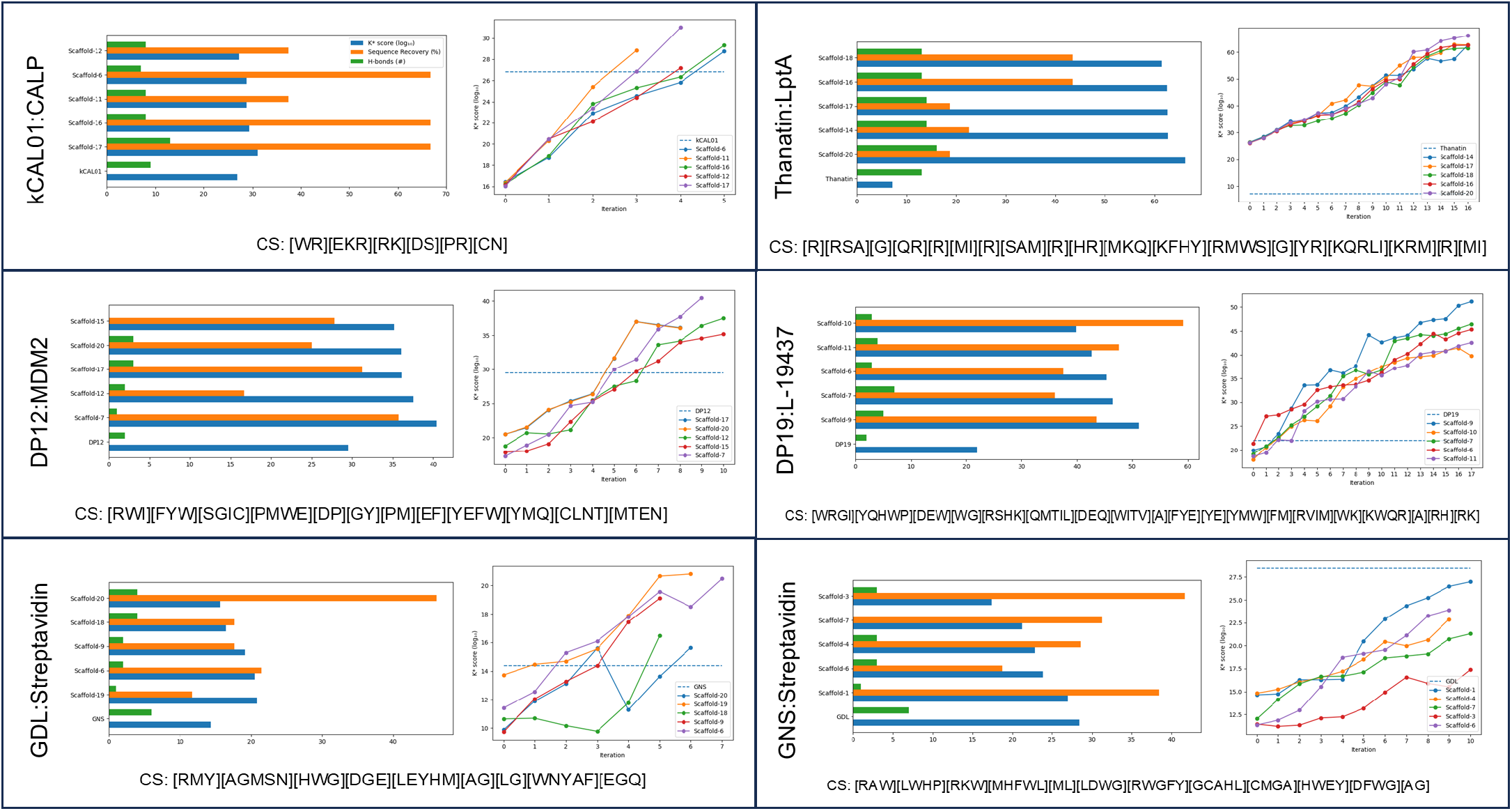
CCK* designs peptides for homochiral and heterochiral systems that converge to high-affinity sequences. The first row shows chirality-preserving homochiral systems, the second row shows chirality-preserving heterochiral systems, and the third row shows chirality-inverting designs. The name of the input template used for design is listed on the left in each cell. **Bar plots:** *K*\* scores (log_10_, blue), sequence recovery (%, orange), and number of hydrogen bonds (green) in the GMEC structure for each prediction. Chirality-inverting designs (GNS, GDL) were compared to the corresponding experimentally solved complex in the opposite chirality (GDL and GNS, respectively). The consensus sequence (CS) is listed below the bar plot with square brackets denoting selected residue identities at each residue index. For example, for L-peptide designs targeting CALP (top left cell), position 1 has identities W and R, position 2 has identities E, K, and R, and so forth. Across these six systems, the consensus sequence degeneracy (number of unique amino acid identities for a given residue position) ranged from 1 to 5 with a mode of 2. **Line plots:** *K*\* scores (log_10_) from the arise algorithm as a function of iteration. Scaffold color labels are listed in the respective legends. The *K*\* score of the known binder is represented as a dashed blue line. For all systems and scaffolds except GNS, arise selects sequences that are predicted to have a better binding affinity than the known peptide binder. Across all arise iterations, the average change in *K*\* score (log_10_) is +1.9 (corresponding to ΔΔ*G* = *−*2.6 kcal/mol).

### Chirality-Preserving Designs

The homochiral systems were kCAL01:CALP (PDB ID 6ov7) and Thanatin:LptA (PDB ID 8gaj), structures solved by our lab using X-ray crystallography. kCAL01 is a 6-residue L-peptide inhibitor that binds the PDZ domain of the CFTR-associated ligand (CALP) in humans, thereby blocking its interaction with CFTR and preventing its CALP-mediated degradation (54). *Podisus maculiventris* Thanatin is a 19-residue L-peptide that binds to the N-terminal *β*-strand of the periplasmic protein LptA, disabling lipopolysaccharide export in Gram-negative bacteria (55). The heterochiral systems were DP12:MDM2 (PDB ID 3iwy) and DP19:L-19437 (PDB ID 7yh8). DP12 is a rationally designed D-peptide antagonist that targets the p53-binding domain of MDM2, representing a promising class of antitumor agents (49). DP19 is a synthetic D-peptide binder selected against the artificial protein target L-19437 (56).

### Chirality-Inverting Designs

The complex Gdlwqheatwkkq:St reptavidin (PDB ID 5n8t, herein referred to as GDL:Streptavidin) is a 9-residue D-peptide bound to L-Streptavidin. The complex GN SFDDWLASKG:Streptavidin (PDB ID 5n89, herein referred to as GNS:Streptavidin) is a 12-residue L-peptide bound to L-Streptavidin. Both of these peptides were designed using a combined stepwise evolution and rational peptide array method (57). GDL and GNS bind Streptavidin with a similar motif and binding site location (complex *C*_α_ alignment RMSD of 0.39 Å), albeit with opposite chirality. Because chirality inversion alters the stereochemical frame of reference, the predicted sequences cannot be benchmarked against the original template. Instead, we initialized the design from the PDB in one chiral space and evaluated the resulting structure against the corresponding experimentally solved complex in the opposite chirality. That is, designs generated using GDL:Streptavidin are benchmarked against GNS:Streptavidin, and vice versa.

For five out of six systems (25 out of 30 designs), sequences designed by *CCK* * were predicted to have a better binding affinity to the target than the corresponding experimentally characterized peptide. Additionally, the consensus sequence degeneracy (number of unique amino acid identities for a given residue position) ranged from 1 to 5 (mode of 2); *CCK* * can therefore converge to high-affinity sequences with low degeneracy. Across all arise iterations, each identity assignment (Eq. 6) produced a mean change in *K*\* score of +1.9 (ΔΔ*G* = −2.6 kcal/mol); this corresponds to an ~80-fold increase in predicted binding affinity. Thus, arise generates sequences that introduce energetically significant improvements comparable in magnitude to a strong buried hydrogen bond or multiple favorable van der Waals contacts. Supplementary data are available in SI Section 2.

## Discussion

We present a computational framework for *de novo* peptide design that operates across both chiral spaces. With three new protein design algorithms (scope, montage, arise), we introduce an approach coupling geometric computation with physics-based sequence evaluation. Previous platforms supporting heterochiral design (10) scaled poorly with the size of the combinatorial sequence space. Despite sampling only a small portion of the theoretical sequence space, *CCK* * converges to sequences with strong predicted binding affinities. These results suggest that *CCK* * likely concentrates search in the productive search space, effectively reducing the exponential design problem for the systems tested here. Because the framework is energy-function agnostic and compatible with any affinity-based search, it can be integrated into existing modeling workflows.

Jain et al. (58) analyze protein design using sparse residue interaction graphs, in which distance-based cutoffs remove edges from the full pairwise interaction graph to reduce computational complexity. *CCK* * is similarly grounded in the observation that much of the difficulty of protein design arises from the combinatorial explosion of pairwise residue couplings. Therefore, these approaches share theoretical motivation: both seek lower-complexity representations of the design landscape that preserve the interactions most relevant to optimality. Instead of reducing complexity by pruning the interaction graph itself, *CCK* * preserves the full *K*\* pairwise energy model by using scope as a fast geometric surrogate for pruning sequence and conformation space before expensive physics-based evaluation.

As previously reported (10), *K*\*-based peptide designs operate within a replication-restitution framework, in which sequences may either reproduce native contacts or adopt alternative favorable interactions. The arise algorithm selects residues that introduce substantial energetic gains (on the order of a strong buried hydrogen bond) while simultaneously avoiding steric clashes (see Fig. 4). Consequently, some designed peptides do not preserve all hydrogen bonds because these contacts are energetically compensated for by alternative favorable interactions (such as multiple favorable van der Waals contacts). Supplementary analyses across independent arise runs suggest that the algorithm is not susceptible to becoming trapped in suboptimal local minima (see SI Section 1).

Future work includes eliminating the requirement for ligand structural inputs to montage, thereby enabling binder design from a target structure alone. Incorporating greater backbone flexibility (15, 59) may further improve design accuracy. Experimental validation is also a valuable future endeavor.

## Source code availability

*CCK* * is available at github.com/donaldlab/OSPREY3.

## Acknowledgments

We received funding from the NIH grant R35 GM-144042 to BRD. We thank all members of the Donald lab for helpful discussions.

## Supplementary Information

### 1 ARISE Local Minima Analysis

While arise employs a greedy search strategy, this approach is not guaranteed to identify the global optimum. Early, locally optimal residue selections could potentially constrain the algorithm to suboptimal solutions. To assess the susceptibility of arise to such local minima, we conducted a series of arise experiments on the kCAL01:CALP system (PDB ID 6ov7), modifying initial residue selections to better evaluate the diversity of local minima found. Although the algorithm does not always find a global minimum, arise finds similarly promising solutions for all initial residue selections; the consensus sequence was [WR][KME]KS[RM]C with a mean *K*\* score of 28.5 ± 1.6.

The highest-affinity alanine scaffold from montage (scaffold-6, *K*\* 16.4) was selected for design. All intersecting mutant convex hulls for peptide residue pairs were computed, yielding (by residue index) the pairs (1, 2), (1, 3), (2, 4), (3, 5), and (4, 6). An arise search was subsequently initiated from each of these pairs. arise designed sequences with *K*\* scores ranging from 25.68 (for the search initiated from (1, 3)) to 30.54 (for the search initiated from (3, 5)), with a mean of 28.5. kCAL01:CALP has a *K*\* score of 26.82; four of five searches yielded sequences with a higher predicted binding affinity (*K*\* score) than the wildtype peptide. See Fig. S1 for a visualization of each arise search.

**Figure S1.**
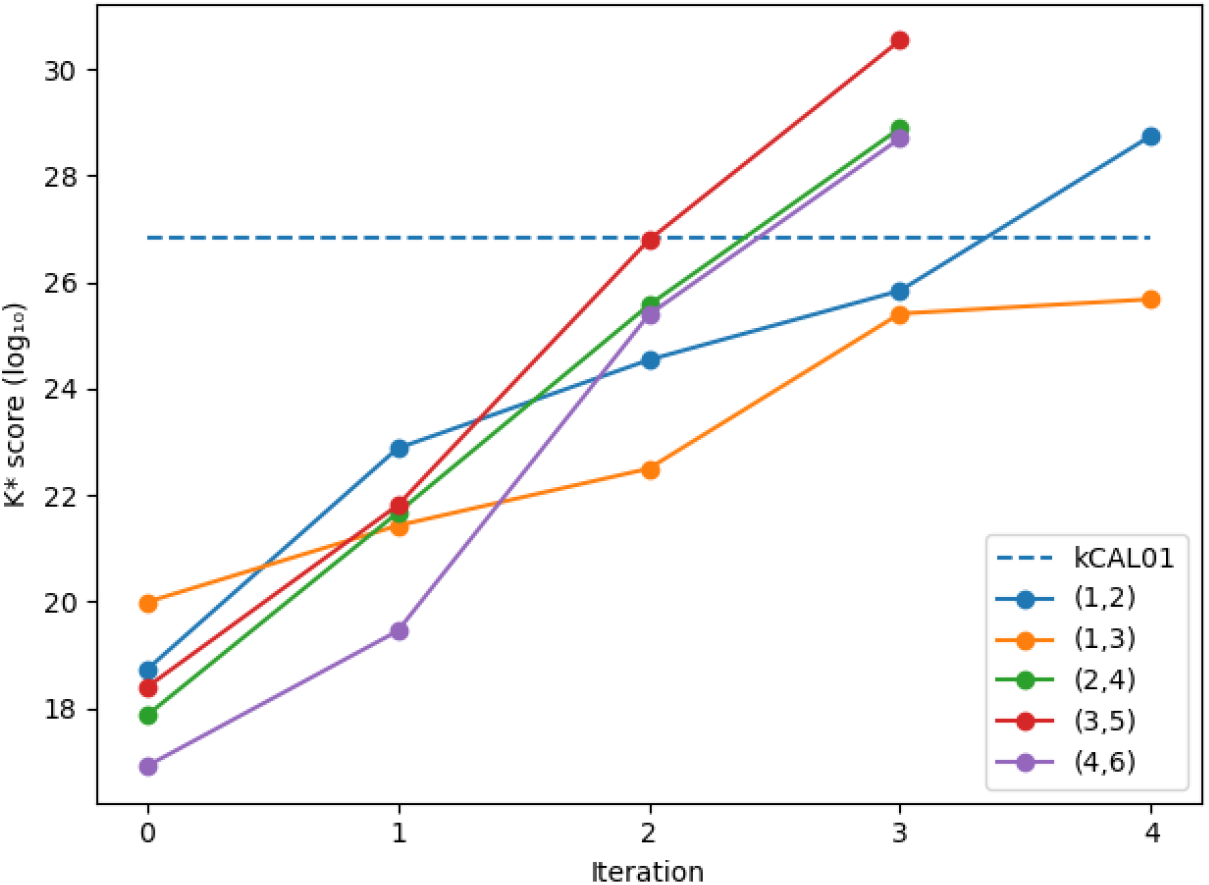
ARISE finds similarly promising solutions for all initial residue selections. *K*\* scores (log_10_) from the arise algorithm as a function of iteration. The *K*\* score of kCAL01:CALP is represented as a dashed blue line. Colors corresponding to each initial residue pair are listed in the legend (e.g., (1, 2) is a search initiated from peptide residue indices 1 and 2).

### 2 Supplementary Results

#### kCAL01:CALP

For kCAL01:CALP, the MASTER subroutine was queried with the 6-residue kCAL01 ligand (WQVTRV), which adopts a linear motif when bound to CALP [1]. The backbone RMSD of the top 20 MASTER returns ranged from 0.05 Å to 0.07 Å, with a median of 0.07 Å. The lowest-RMSD match was the endogenous ligand kCAL01 (RMSD 0 Å), and was therefore discarded. The alanine scaffolds produced by montage resulted in *K*\* scores ranging from 6.65 (scaffold-4) to 16.42 (scaffold-6), with a median of 15.81. The five highest-affinity alanine scaffolds returned by montage were scaffold-6 (RMSD 0.06 Å, *K*\* 16.42), scaffold-11 (RMSD 0.07 Å, *K*\* 16.37), scaffold-16 (RMSD 0.07 Å, *K*\* 16.22), scaffold-12 (RMSD 0.07 Å, *K*\* 16.14), and scaffold-17 (RMSD 0.07 Å, *K*\* 16.02).

The arise algorithm enumerated a total of 2,988 sequences across these five backbones. The number of enumerated sequences ranged from 452 (scaffold-12) to 819 (scaffold-17), with a median of 459. The *K*\* search runtime for all iterations ranged from 21 minutes (scaffold-6) to 1,467 minutes (scaffold-12), with a median runtime of 74 minutes (scaffold-11). Across all arise iterations, the average change in *K*\* score is +3.15. Full-sequence *K*\* scores ranged from 27.19 (scaffold-12) to 31.02 (scaffold-17), with a median score of 28.84 (scaffold-11). Scaffold-17 produced the tightest binder (*K*\* 31.02) with sequence WERSRN. kCAL01:CALP has a *K*\* score of 26.82; therefore, all designed sequences have a higher predicted binding affinity for CALP than kCAL01.

The consensus sequence across the five backbones is [WR][EKR][RK][DS][PR][CN]. The sequence similarity of highest-affinity scaffold-17 (WERSRN) to kCAL01 (WQVTRV) is 66.67%. The sequence similarity ranged from 37.50% (scaffold-11, 12) to 66.67% (scaffold-6, 16, 17), with a median of 66.67%. The consensus sequence degeneracy ranged from 2 to 3 (mode of 2). kCAL01:CALP has 9 H-bonds. The change in the number of H-bonds ranges from −1 (scaffold-11, 12, 16) to +4 (scaffold-17), with a median change of −1 hydrogen bond.

#### Thanatin:LptA

For Thanatin:LptA, the MASTER subroutine was queried with the 19-residue Thanatin ligand (KKPVPIIYCNRRTGKCQRM), which adopts a *β*-hairpin motif when bound to LptA [2]. The backbone RMSD of the top 20 MASTER returns ranged from 0.75 Å to 0.99 Å, with a median of 0.92 Å. The alanine scaffolds produced by montage resulted in *K*\* scores ranging from 19.06 (scaffold-5) to 23.42 (scaffold-14), with a median *K*\* score of 22.29. The five highest-affinity alanine scaffolds returned by montage were scaffold-14 (RMSD 0.98 Å, *K*\* 23.42), scaffold-12 (RMSD 0.96 Å, *K*\* 23.05), scaffold-17 (RMSD 0.99 Å, *K*\* 22.87), scaffold-18 (RMSD 0.99 Å, *K*\* 22.79), and scaffold-16 (RMSD 0.98 Å, *K*\* 22.75). Scaffold-12 exhibited a technical failure during preprocessing, preventing arise from completing the design run. To maintain a consistent sample size across backbones, we substituted with the next-highest-ranked backbone (scaffold-20: RMSD 0.99 Å, *K*\* 22.72).

The arise algorithm enumerated a total of 1,690 sequences across these five backbones. Each backbone enumerated 338 sequences across all iterations. The *K*\* search runtime for all iterations ranged from 81 minutes (scaffold-16) to 1,809 minutes (scaffold-17), with a median runtime of 235 minutes (scaffold-14). Across all arise iterations, the average change in *K*\* score is +2.30. Full-sequence *K*\* scores ranged from 61.47 (scaffold-18) to 66.15 (scaffold-20), with a median score of 62.65 (scaffold-17). Scaffold-20 produced the tightest binder (*K*\* 66.15) with sequence RRGQRMRSRHMKRGYKKRM. Thanatin:LptA has a *K*\* score of 7.15; therefore, all designed sequences have a higher predicted binding affinity for LptA than Thanatin.

The consensus sequence across the five backbones is [R][RSA][G][QR][R][MI][R][SAM][R][HR][MKQ][KF HY][RMWS][G][YR][KQRLI][KRM][R][MI]. The sequence similarity of highest-affinity scaffold-20 (RRGQR MRSRHMKRGYKKRM) to Thanatin (KKPVPIIYCNRRTGKCQRM) is 18.75%. The sequence similarity ranged from 18.75% (scaffold-17, 20) to 43.48% (scaffold-16, 18), with a median of 22.58% (scaffold-14). The consensus sequence degeneracy ranged from 1 to 5 (mode of 1). Thanatin:LptA has 13 H-bonds. The change in the number of H-bonds ranges from 0 (scaffold-16, 18) to +3 (scaffold-20), with a median of +1 (scaffold-14, 17). No designs reduced the total number of hydrogen bonds relative to Thanatin:LptA.

#### DP12:MDM2

For DP12:MDM2, the MASTER subroutine was queried with the 12-residue DP12 ligand (DWWPLAFEALLR), which adopts an *α*-helix motif when bound to MDM2. The backbone RMSD of the top 20 MASTER returns ranged from 0.43 Å to 0.49 Å, with a median of 0.46 Å. The alanine scaffolds produced by montage resulted in *K*\* scores ranging from 12.42 (scaffold-5) to 17.55 (scaffold-17), with a median *K*\* score of 15.20. The five highest-affinity alanine scaffolds returned by montage were scaffold-17 (RMSD 0.48 Å, *K*\* 17.55), scaffold-20 (RMSD 0.49 Å, *K*\* 17.53), scaffold-12 (RMSD 0.46 Å, *K*\* 16.43), scaffold-15 (RMSD 0.47 Å, *K*\* 16.24), and scaffold-7 (RMSD 0.45 Å, *K*\* 15.83).

The arise algorithm enumerated a total of 1,351 sequences across these five backbones. The number of enumerated sequences ranged from 178 (scaffold-17, 20) to 579 (scaffold-12), with a median of 198 (scaffold-7). The *K*\* search runtime for all iterations ranged from 7 minutes (scaffold-15) to 169 minutes (scaffold-12), with a median runtime of 9 minutes (scaffold-7). Across all arise iterations, the average change in *K*\* score is +2.01. Full-sequence *K*\* scores ranged from 35.17 (scaffold-15) to 40.43 (scaffold-7), with a median score of 36.11 (scaffold-17). Scaffold-7 produced the tightest binder (*K*\* 40.43) with sequence WYGMPYMEEMCT. DP12:MDM2 has a *K*\* score of 29.49; therefore, all designed sequences have a higher predicted binding affinity for MDM2 than DP12.

The consensus sequence across the five backbones is [RWI][FYW][SGIC][PMWE][DP][GY][PM][EF][Y EFW][YMQ][CLNT][MTEN]. The sequence similarity of highest-affinity scaffold-7 (WYGMPYMEEMCT) to DP12 (DWWPLAFEALLR) is 35.71%. The sequence similarity ranged from 16.67% (scaffold-12) to 35.71% (scaffold-7), with a median of 27.78% (scaffold-15). The consensus sequence degeneracy ranged from 2 to 4 (mode of 4). DP12:MDM2 has 2 H-bonds. The change in the number of H-bonds ranges from −2 (scaffold-15) to +1 (scaffold-17, 20), with a median of 0 (scaffold-12).

#### DP19:L-19437

For DP19:L-19437, the MASTER subroutine was queried with the 19-residue DP19 ligand (DEHELLETAARWFYEIAKR), which adopts an *α*-helix motif when bound to L-19437. The backbone RMSD of the top 20 MASTER returns ranged from 0.18 Å to 0.20 Å, with a median of 0.20 Å. The alanine scaffolds produced by montage resulted in *K*\* scores ranging from 15.70 (scaffold-15) to 18.73 (scaffold-9), with a median *K*\* score of 16.72. The five highest-affinity alanine scaffolds returned by montage were scaffold-9 (RMSD 0.20 Å, *K*\* 18.73), scaffold-10 (RMSD 0.20 Å, *K*\* 17.23), scaffold-7 (RMSD 0.19 Å, *K*\* 17.19), scaffold-6 (RMSD 0.19 Å, *K*\* 17.19), and scaffold-11 (RMSD 0.20 Å, *K*\* 17.11).

The arise algorithm enumerated a total of 2,152 sequences across these five backbones. The number of enumerated sequences ranged from 358 (scaffold-6, 9, 11) to 719 (scaffold-7), with a median of 358. Across all arise iterations, the average change in *K*\* score is +1.51. The *K*\* search runtime for all iterations ranged from 11 minutes (scaffold-11) to 28 minutes (scaffold-6), with a median runtime of 16 minutes (scaffold-9). Full-sequence *K*\* scores ranged from 39.78 (scaffold-10) to 51.17 (scaffold-9), with a median score of 45.34 (scaffold-6). Scaffold-9 produced the tightest binder (*K*\* 51.17) with sequence RHEGHTEWAYEMFVK WARR. DP19:L-19437 has a *K*\* score of 21.91; therefore, all designed sequences have a higher predicted binding affinity for L-19437 than DP19.

The consensus sequence across the five backbones is [WRGI][YQHWP][DEW][WG][RSHK][QMTIL][DE Q][WITV][A][FYE][YE][YMW][FM][RVIM][WK][KWQR][A][RH][RK]. The sequence similarity of highest-affinity scaffold-9 (RHEGHTEWAYEMFVKWARR) to DP19 (DEHELLETAARWFYEIAKR) is 43.48%. The sequence similarity ranged from 36.00% (scaffold-7) to 59.09% (scaffold-10), with a median of 43.48% (scaffold-9). The consensus sequence degeneracy ranged from 1 to 5 (mode of 2). DP19:L-19437 has 2 H-bonds. The change in the number of H-bonds ranges from +1 (scaffold-6, 10) to +5 (scaffold-7), with a median of +2 (scaffold-11). All DP19 designs are predicted to establish at least one additional hydrogen bond with L-19437.

#### GDL:Streptavidin

The heterochiral system GDL:Streptavidin was used to design L-peptide binders. The MASTER subroutine was queried with the GDL ligand (LWQHEATWK), which adopts an *α*-helix motif when bound to Streptavidin. The backbone RMSD of the top 20 MASTER returns ranged from 1.21 Å to 1.53 Å, with a median of 1.48 Å. The alanine scaffolds produced by montage resulted in *K*\* scores ranging from 4.89 (scaffold-14) to 9.98 (scaffold-20), with a median *K*\* score of 7.63. The five highest-affinity alanine scaffolds returned by montage were scaffold-20 (RMSD 1.53 Å, *K*\* 9.98), scaffold-19 (RMSD 1.52 Å, *K*\* 9.18), scaffold-18 (RMSD 1.52 Å, *K*\* 9.02), scaffold-9 (RMSD 1.48 Å, *K*\* 8.80), and scaffold-6 (RMSD 1.35 Å, *K*\* 8.21).

The arise algorithm enumerated a total of 1,026 sequences across these five backbones. The number of enumerated sequences ranged from 118 (scaffold-9, 18) to 519 (scaffold-6), with a median of 133 (scaffold-20). Across all arise iterations, the average change in *K*\* score is +1.30. The *K*\* search runtime for all iterations ranged from 6 minutes (scaffold-9) to 2,165 minutes (scaffold-20), with a median runtime of 11 minutes (scaffold-19). Full-sequence *K*\* scores ranged from 15.64 (scaffold-20) to 20.80 (scaffold-19), with a median score of 19.10 (scaffold-9). Scaffold-19 produced the tightest binder (*K*\* 20.80) with sequence RM WEYGGYQ. GNS:Streptavidin (corresponding experimental structure) has a *K*\* score of 14.38; therefore, all designed sequences have a higher predicted binding affinity for Streptavidin than GNS.

The consensus sequence across the five backbones is [RMY][AGMSN][HWG][DGE][LEYHM][AG][LG][W NYAF][EGQ]. The sequence similarity of highest-affinity scaffold-19 (RMWEYGGYQ) to GNS (GNSFDDWLASKG) is 11.76%. The sequence similarity ranged from 11.76% (scaffold-19) to 46.15% (scaffold-20), with a median of 17.65%. The consensus sequence degeneracy ranged from 2 to 5 (mode of 3). GNS:Streptavidin (corresponding experimental structure) has 6 H-bonds. The change in the number of H-bonds ranges from *−*5 (scaffold-19) to *−*2 (scaffold-18, 20), with a median of *−*4 (scaffold-6, 9).

#### GNS:Streptavidin

The homochiral system GNS:Streptavidin was used to design D-peptide binders. For GNS:Streptavidin, the MASTER subroutine was queried with GNS (GNSFDDWLASKG), which adopts an *α*-helix motif when bound to Streptavidin. The backbone RMSD of the top 20 MASTER returns ranged from 1.11 Å to 2.31 Å, with a median of 2.31 Å. The alanine scaffolds produced by montage resulted in *K*\* scores ranging from 8.77 (scaffold-2) to 13.90 (scaffold-1), with a median *K*\* score of 9.62. The five highest-affinity alanine scaffolds returned by montage were scaffold-1 (RMSD 1.11 Å, *K*\* 13.90), scaffold-4 (RMSD 2.00 Å, *K*\* 12.07), scaffold-7 (RMSD 2.15 Å, *K*\* 10.79), scaffold-3 (RMSD 1.89 Å, *K*\* 10.48), and scaffold-6 (RMSD 2.12 Å, *K*\* 10.07).

The arise algorithm enumerated a total of 1,394 sequences across these five backbones. The number of enumerated sequences ranged from 192 (scaffold-4) to 579 (scaffold-1), with a median of 212 (scaffold-7). Across all arise iterations, the average change in *K*\* score is +1.01. The *K*\* search runtime for all iterations ranged from 56 minutes (scaffold-6) to 1,530 minutes (scaffold-4), with a median runtime of 1,465 minutes (scaffold-7). Full-sequence *K*\* scores ranged from 17.39 (scaffold-3) to 26.99 (scaffold-1), with a median score of 22.88 (scaffold-4). Scaffold-1 produced the tightest binder (*K*\* 26.99) with sequence APRLMW YLAYWA. GDL:Streptavidin (corresponding experimental structure) has a *K*\* score of 28.43. Thus, the highest-scoring design did not exceed the predicted binding affinity of the corresponding GDL:Streptavidin experimental structure.

The consensus sequence across the five backbones is [RAW][LWHP][RKW][MHFWL][ML][LDWG][RWG FY][GCAHL][CMGA][HWEY][DFWG][AG]. The sequence similarity of highest-affinity scaffold-1 (APRLM WYLAYWA) to GDL (LWQHEATWK) is 38.46%. The sequence similarity ranged from 18.75% (scaffold-6) to 41.67% (scaffold-3), with a median of 31.25% (scaffold-7). The consensus sequence degeneracy ranged from 2 to 5 (mode of 4). GDL:Streptavidin (corresponding experimental structure) has 7 H-bonds. The change in the number of H-bonds ranges from *−*7 (scaffold-7) to *−*4 (scaffold-3, 4, 6), with a median of *−*4.

**Table S1.**
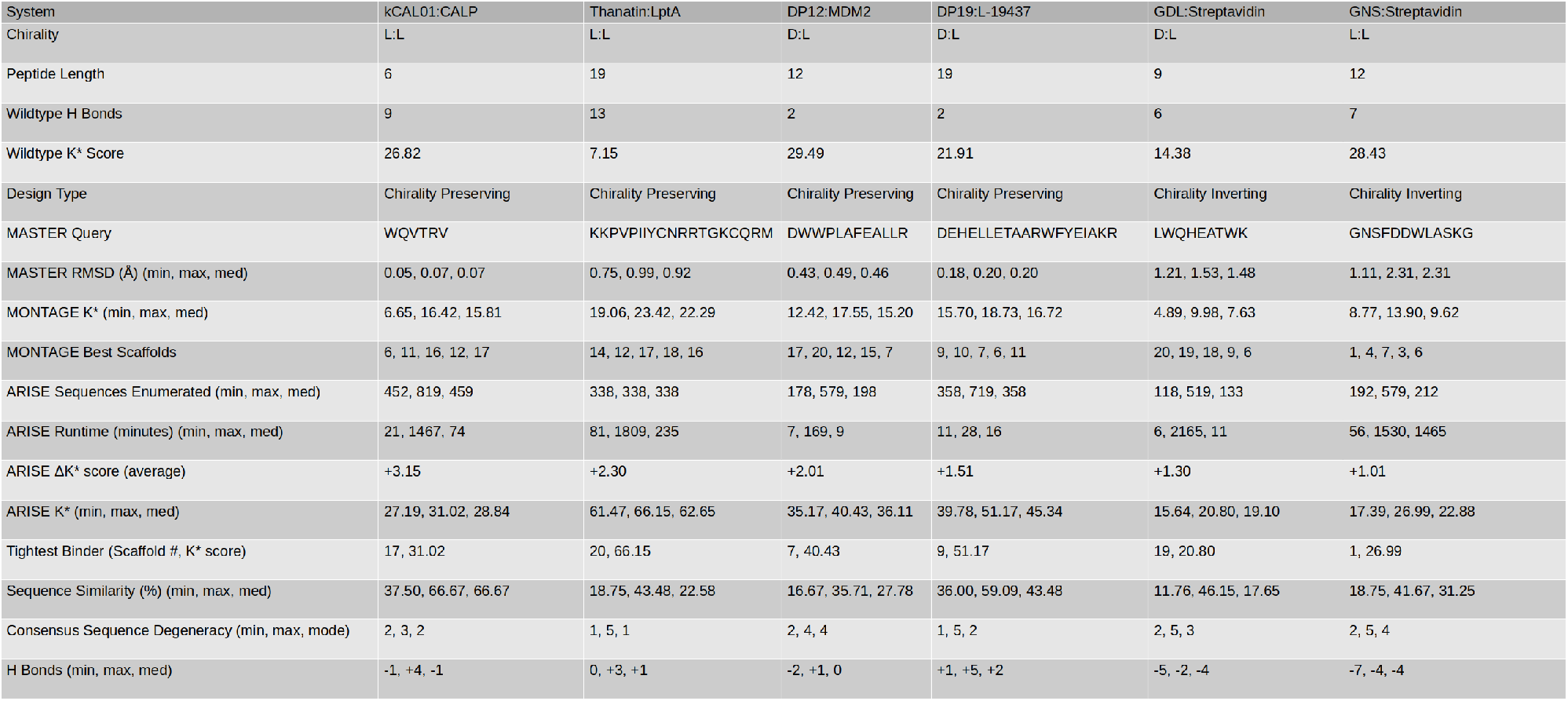
Summary Statistics. All *K*\* scores are on a log_10_ scale. Chirality reports the chirality of the peptide:target complex. Peptide Length reports the length of the peptide. Wildtype H Bonds reports the number of hydrogen bonds in the wildtype experimental structure, and Wildtype *K*\* Score reports the *K*\* score of the experimental structure. Design Type indicates chirality-preserving (e.g., L to L) and chirality-inverting (e.g., L to D) designs. MASTER Query is the sequence of the input structure to the MASTER subroutine in the montage algorithm. MASTER RMSD (Å) (min, max, med) reports the minimum, maximum, and median RMSD of MASTER matches in angstroms. montage *K*\* (min, max, med) reports the minimum, maximum, and median *K*\* score for alanine scaffolds produced by montage. montage Best Scaffolds reports the five highest-affinity scaffolds produced by montage, ordered by decreasing *K*\* score. arise Sequences Enumerated (min, max, med) reports the minimum, maximum, and median number of sequences enumerated across all arise steps for a given scaffold input. arise Runtime (minutes) (min, max, med) reports the minimum, maximum, and median runtime in minutes across all arise steps for a given scaffold input. arise Δ*K*\* score (average) reports the average change in *K*\* score across all arise iterations. arise *K*\* (min, max, med) is the minimum, maximum, and median *K*\* score across all designed sequences. Tightest Binder (Scaffold #, *K*\* score) reports the scaffold number (ordered by MASTER RMSD) and *K*\* score of the highest-affinity design. Sequence Similarity (%) (min, max, med) reports the minimum, maximum, and median sequence similarity across all designs relative to the corresponding peptide sequence in the experimental structure. Consensus Sequence Degeneracy (min, max, mode) reports the minimum, maximum, and mode of the consensus sequence degeneracy (number of unique amino acid identities for a given residue position). H Bonds (min, max, med) reports the minimum, maximum, and median number of hydrogen bonds in the sequences designed by arise relative to the corresponding experimental structure.

**Figure S2.**
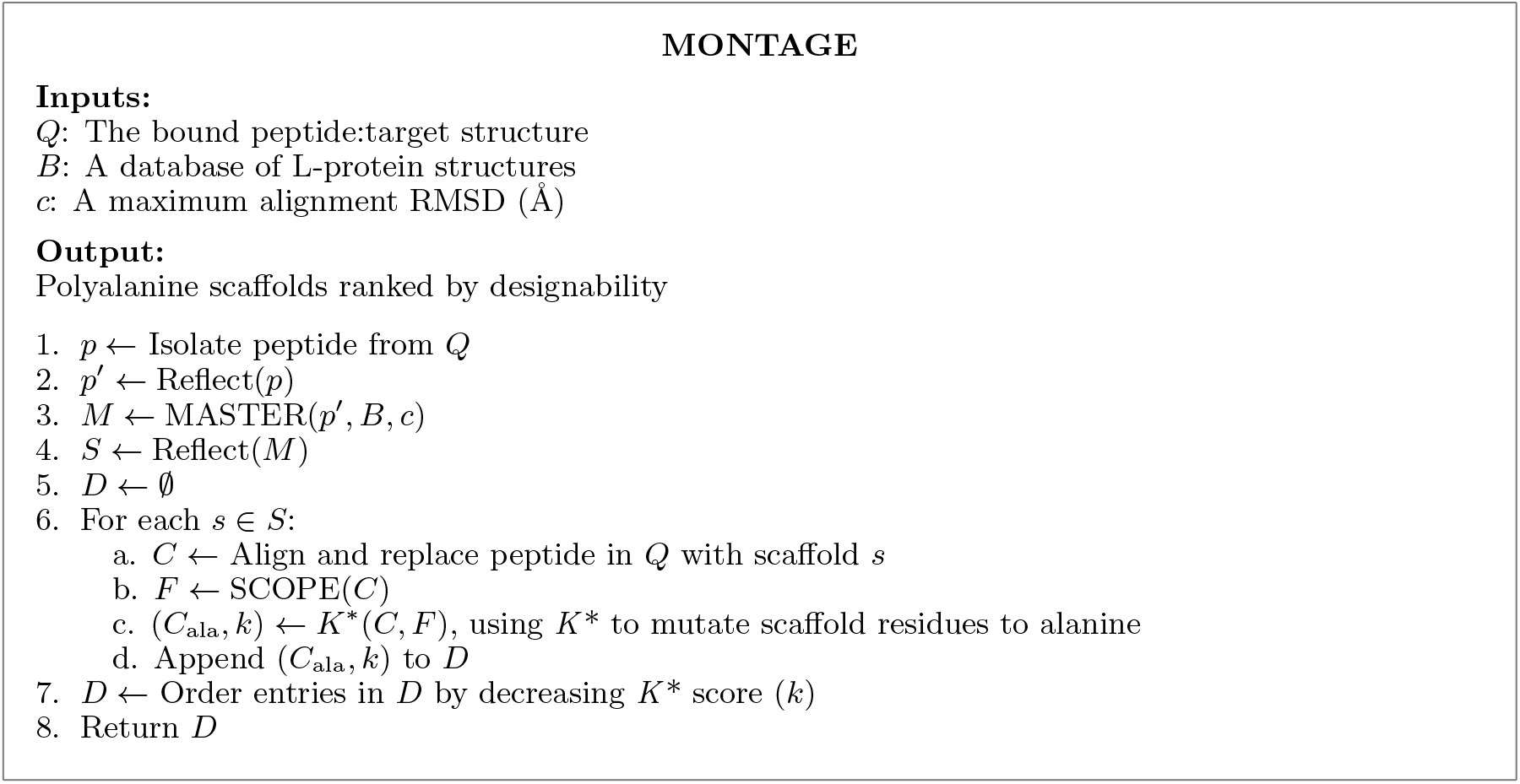
MONTAGE pseudocode for D-peptide scaffold generation. montage generates D-peptide scaffold libraries by isolating a template L-peptide, reflecting it, and using MASTER [3] to identify structurally similar backbones within a *C*_*α*_ alignment RMSD cutoff. MASTER returns backbone matches in order of increasing RMSD. Backbones are reflected once more to generate D-peptide matches that mimic the L-peptide template. D-scaffolds are aligned into the target binding interface to replace the template L-peptide. The *K*\* algorithm (with sampling parameterized by scope) mutates scaffold residues to alanine and ranks by predicted binding affinity. Together, *C*_*α*_ alignment RMSD and *K*\* score determine scaffold designability: MASTER identifies and orders candidate backbones by RMSD, while *K*\* ranks the polyalanine scaffolds by affinity. L-peptide scaffolds are generated by ablating reflection operations.

## References

1. Lei Wang, Nanxi Wang, Wenping Zhang et al. Therapeutic peptides: current applications and future directions. Signal Transduction and Targeted Therapy, 2022.

2. Li Di. Strategic approaches to optimizing peptide ADME properties. The AAPS Journal, 2015.

3. Niles A. Pierce and Erik Winfree. Protein design is NP-hard. Protein Engineering, Design, and Selection, 2002.

4. Pernille Vosbein, Paula Paredes Vergara, Danny T Huang et al. AlphaFold ensemble competition screens enable peptide binder design with single-residue sensitivity. ACS Chemical Biology, 2024.

5. Qiuzhen Li, Efstathios Nikolaos Vlachos, and Patrick Bryant. Design of linear and cyclic peptide binders from protein sequence information. Communications Chemistry, 2025.

6. Qiuzhen Li, Diandra Daumiller, and Patrick Bryant. RareFold: Structure prediction and design of proteins with noncanonical amino acids. bioRxiv, 2025.

7. Jin Sub Lee and Philip M. Kim. Design of peptides with non-canonical amino acids using flow matching. bioRxiv, 2025.

8. Henry Childs, Pei Zhou, and Bruce R. Donald. AlphaFold 3 fails to predict D-peptide chirality, fold, and binding pose in heterochiral complexes. bioRxiv, 2026.

9. Michael Garton, Satra Nim, Tracy A. Stone et al. Method to generate highly stable D-amino acid analogs of bioactive helical peptides using a mirror image of the entire PDB. Proc. Natl. Acad. Sci. U.S.A., 2018.

10. Nathan Guerin, Henry Childs, Pei Zhou et al. DexDesign: an OSPREY-based algorithm for designing de novo D-peptide inhibitors. Protein Engineering, Design and Selection, 2024.

11. Henry Childs, Nathan Guerin, Pei Zhou et al. Protocol for designing de novo noncanonical peptide binders in OSPREY. J Comput. Biol., 2024.

12. Karoline Santur, Elke Reinartz, Yi Lien et al. Discovery of all-D-peptide inhibitors of SARS-CoV-2 3C-like protease. ACS Chemical Biology, 2023.

13. T. Schumacher, L. Mayr, D. Minor Jr et al. Identification of D-peptide ligands through mirror-image phage display. Science, 1996.

14. Bassil I. Dahiyat, Catherine A. Sarisky, and Stephen L. Mayo. De novo protein design: Towards fully automated sequence selection. Science, 1997.

15. Mark A. Hallen, Daniel A. Keedy, and Bruce R. Donald. Dead-end elimination with perturbations (DEEPer): A provable protein design algorithm with continuous sidechain and backbone flexibility. Proteins, 2012.

16. Graham T Holt, Jason Gorman, Siyu Wang et al. Improved HIV-1 neutralization breadth and potency of V2-apex antibodies by in silico design. Cell Reports, 2023.

17. Anna U. Lowegard, Marcel S. Frenkel, Graham T. Holt et al. Novel, provable algorithms for efficient ensemble-based computational protein design and their application to the redesign of the c-Raf-RBD:KRas protein-protein interface. PLOS Computational Biology, 2020.

18. Siyu Wang, Stephanie M. Reeve, Graham T. Holt et al. Chiral evasion and stereospecific antifolate resistance in Staphylococcus aureus. PLOS Computational Biology, 2022.

19. PD Renfrew, EJ Choi, R Bonneau et al. Incorporation of noncanonical amino acids into Rosetta and use in computational protein-peptide interface design. PLoS ONE, 2012.

20. G Bhardwaj, VK Mulligan, and CD Bahl. Accurate de novo design of hyperstable constrained peptides. Nature, 2016.

21. Seydou Traoré, David Allouche, Isabelle André et al. A new framework for computational protein design through cost function network optimization. Bioinformatics, 2013.

22. Daniel V. Schroeder. An Introduction to Thermal Physics. Oxford University Press, 2021.

23. Rommie E. Amaro, Jerome Baudry, John Chodera et al. Ensemble docking in drug discovery. Biophysical Journal, 2018.

24. Jeremy Wohlwend, Gabriele Corso, Saro Passaro et al. Boltz-1 democratizing biomolecular interaction modeling. bioRxiv, 2025.

25. R. H. Lilien, B. W. Stevens, A. C. Anderson et al. A novel ensemble-based scoring and search algorithm for protein redesign and its application to modify the substrate specificity of the gramicidin synthetase A phenylalanine adenylation enzyme. Journal of Computational Biology, 12:740–761, 2005.

26. Pablo Gainza, Kyle E. Roberts, and Bruce R. Donald. Protein design using continuous rotamers. PLOS Computational Biology, 2012.

27. Mark A Hallen, Jeffrey W Martin, Adegoke Ojewole et al. OSPREY 3.0: Open-source protein redesign for you, with powerful new features. J Comput Chem, 2018.

28. A. A. Ojewole, J. D. Jou, V. G. Fowler et al. BBK* (Branch and Bound over K*): A provable and efficient ensemble-based algorithm to optimize stability and binding affinity over large sequence spaces. Journal of Computational Biology, 25:726–739, 2018.

29. Jonathan D Jou, Graham T Holt, Anna U Lowegard et al. Minimization-Aware Recursive K*: A novel, provable algorithm that accelerates ensemble-based protein design and provably approximates the energy landscape. J Comput Biol, 2020.

30. Ivelin Georgiev, Ryan H. Lilien, and Bruce R. Donald. The minimized dead-end elimination criterion and its application to protein redesign in a hybrid scoring and search algorithm for computing partition functions over molecular ensembles. J. Comput. Chem., 2008.

31. David A. Case, David S. Cerutti, Vinícius Wilian D. Cruzeiro et al. Recent developments in Amber biomolecular simulations. J. Chem. Inf. Model., 2025.

32. Themis Lazaridis and Martin Karplus. Effective energy function for proteins in solution. Proteins, 1999.

33. Andrew R. Leach and Andrew P. Lemon. Exploring the conformational space of protein side chains using dead-end elimination and the A* algorithm. Proteins, 1998.

34. Anna Lowegard. Novel Algorithms and Tools for Computational Protein Design with Applications to Drug Resistance Prediction, Antibody Design, Peptide Inhibitor Design, and Protein Stability Prediction. PhD thesis, Duke University, 2019.

35. Simon C. Lovell, J. Michael Word, Jane S. Richardson et al. The penultimate rotamer library. Proteins, 2000.

36. E. Noether. Gesammelte Abhandlungen-collected papers, Springer collected works in mathematics. Springer, 1983.

37. Zhiyong Wang and Jinbo Xu. Predicting protein contact map using evolutionary and physical constraints by integer programming. Bioinformatics, 2013.

38. Sheng Wang, Siqi Sun, Zhen Li et al. Accurate de novo prediction of protein contact map by ultra-deep learning model. PLoS Comput. Biol., 2017.

39. John Jumper, Richard Evans, Alexander Pritzel et al. Highly accurate protein structure prediction with AlphaFold. Nature, 2021.

40. Mark A. Hallen. PLUG (Pruning of Local Unrealistic Geometries) removes restrictions on biophysical modeling for protein design. Proteins, 2018.

41. Jonathan D Jou, Swati Jain, Ivelin S Georgiev et al. BWM*: A novel, provable, ensemble-based dynamic programming algorithm for sparse approximations of computational protein design. J Comput Biol., 2016.

42. C. Bradford Barber, David P. Dobkin, and Hannu Huhdanpaa. The quickhull algorithm for convex hulls. ACM TOMS, 1996.

43. Bane Sullivan and Alexander Kaszynski. PyVista: 3D plotting and mesh analysis through a streamlined interface for the Visualization Toolkit (VTK). Journal of Open Source Software, 2019.

44. Jianfu Zhou and Gevorg Grigoryan. Rapid search for tertiary fragments reveals protein sequence–structure relationships. Proteins, 2014.

45. Merve Ayyildiz, Jakob Noske, Florian J. Gisdon et al. Evaluation of physics-based protein design methods for predicting single residue effects on peptide binding specificities. J. Comput. Chem, 2025.

46. H.M. Berman, J. Westbrook, Z. Feng et al. The protein data bank. Nucleic Acids Research, 2000.

47. Michel van Kempen, Stephanie S. Kim, Charlotte Tumescheit et al. Fast and accurate protein structure search with foldseek. Nat. Biotechnol., 2024.

48. Hetunandan Kamisetty, Eric P. Xing, and Christopher J. Langmead. Free energy estimates of all-atom protein structures using generalized belief propagation. J. Comput. Biol, 2008.

49. Min Liu, Chong Li, Marzena Pazgier et al. D-peptide inhibitors of the p53–MDM2 interaction for targeted molecular therapy of malignant neoplasms. Proc. Natl. Acad. Sci. U.S.A., 2010.

50. Christopher J Williams, Jeffrey J Headd, Nigel W Moriarty et al. Molprobity: More and better reference data for improved all-atom structure validation. Protein Sci., 2018.

51. Jonathan Jou, Nathan Guerin, and Kyle Roberts. Protein design plugin, accessed 2026.

52. Schrödinger, LLC. The PyMOL molecular graphics system, version 2.5. The PyMOL Molecular Graphics System, Version 2.5, Schrödinger, LLC., accessed 2025.

53. VectorBuilder. Sequence alignment, accessed 2025.

54. Graham T. Holt, Jonathan D. Jou, Nicholas P. Gill et al. Computational analysis of energy landscapes reveals dynamic features that contribute to binding of inhibitors to CFTR-associated ligand. J. Phys. Chem. B., 2019.

55. Kelly Huynh, Amanuel Kibrom, Bruce R. Donald et al. Discovery, characterization, and redesign of potent antimicrobial thanatin orthologs from Chinavia ubica and Murgantia histrionica targeting E. coli LptA. Journal of Structural Biology, 2023.

56. Ke Sun, Sicong Li, Bowen Zheng et al. Accurate de novo design of heterochiral protein–protein interactions. Cell Res, 2024.

57. Victor I. Lyamichev, Lauren E. Goodrich, Eric H. Sullivan et al. Stepwise evolution improves identification of diverse peptides binding to a protein target. Sci Rep, 2017.

58. Swati Jain, Jonathan D. Jou, Ivelin S. Georgiev et al. A critical analysis of computational protein design with sparse residue interaction graphs. PLoS Comput. Biol., 2017.

59. Allen C. McBride, Feng Yu, Edward H. Cheng et al. Predicting pose distribution of protein domains connected by flexible linkers is an unsolved problem. Proteins, 2025.

## References

[1] Nathan Guerin, Henry Childs, Pei Zhou, and Bruce R Donald. “DexDesign: an OSPREY-based algorithm for designing de novo D-peptide inhibitors”. In: Protein Engineering, Design and Selection (2024). doi: 10.1093/protein/gzae007.

[2] Kelly Huynh, Amanuel Kibrom, Bruce R. Donald, and Pei Zhou. “Discovery, characterization, and redesign of potent antimicrobial thanatin orthologs from Chinavia ubica and Murgantia histrionica targeting E. coli LptA”. In: Journal of Structural Biology (2023). doi: 10.1016/j.yjsbx.2023.100091.

[3] Jianfu Zhou and Gevorg Grigoryan. “Rapid search for tertiary fragments reveals protein sequence–structure relationships”. In: Proteins (2014). doi: 10.1002/pro.2610.

